# Classification of helical polymers with deep-learning language models

**DOI:** 10.1101/2023.07.28.550909

**Authors:** Daoyi Li, Wen Jiang

## Abstract

Many macromolecules in biological systems exist in the form of helical polymers. However, the inherent polymorphism and heterogeneity of samples complicate the reconstruction of helical polymers from cryo-EM images. Currently available 2D classification methods are effective at separating particles of interest from contaminants, but they do not effectively differentiate between polymorphs, resulting in heterogeneity in the 2D classes. As such, it is crucial to develop a method that can computationally divide a dataset of polymorphic helical structures into homogenous subsets. In this work, we utilized deep-learning language models to embed the filaments as vectors in hyperspace and group them into clusters. Tests with both simulated and experimental datasets have demonstrated that our method – HLM (**H**elical classification with **L**anguage **M**odel) can effectively distinguish different types of filaments, in the presence of many contaminants and low signal-to-noise ratios. We also demonstrate that HLM can isolate homogeneous subsets of particles from a publicly available dataset, resulting in the discovery of a previously unknown non-proteinaceous density around tau filaments.

## 1. Introduction

Many macromolecules in biological systems exist in the form of helical polymers, such as amyloid fibrils^1–3^, microtubule^4^, actin^5, 6^, bacteria pili^7^, viruses^8^, and phage tails^9^. 3D reconstruction of these helical polymers is essential to elucidate the structure-function relationships and provide an understanding of related diseases at the atomic level^10, 11^, leading to potential drug discovery^12^. Often, the 3D reconstruction of helical polymers is challenged by the heterogeneity of the sample^13^.

Helical reconstruction from cryo-EM images involves the deposition of filaments on grids and the collection of projection image data. Then, the filaments on these images are identified manually or through automated methods. The traditional method of reconstructing the 3D structure of helical polymers is performed in the Fourier space of the full-length filament. The helical parameters are determined through indexing of the Fourier layer lines, and the reconstruction is based on Fourier-Bessel synthesis^9, 14, 15^. However, this method requires a long, straight, and uniform helical structure. When the helical structures are short, poorly ordered, or tilted out of the plane, the layer lines become weak and ambiguous, making it challenging to derive helical parameters using the Fourier-Bessel indexing of helical parameters^16, 17^. Reconstruction using iterative helical real-space reconstruction (IHRSR) does not require the helical polymers to be as long or straight. This is because the helical polymers are computationally segmented along the helical axis, and the segments are analyzed like “single particles” and then reconstructed using the estimated helical parameter^18–20^. However, the presence of heterogeneous types of helical structures can impact the accuracy of the estimated helical parameters, separation of the polymorphs into homogenous subsets, and ultimately affect the final reconstructions^20, 21^.

There are several approaches to separate different types of filaments, including manual selection^1, 3^, 2D classification^22^, and 3D classification^23^. Manual picking requires visually distinct features, for example, diameter or shape, and the researcher must have sufficient prior knowledge of structural differences in the helical assemblies. Distinct types of helical structures can also have a similar diameter or features beyond visual recognition^23^. Manual selection is thus not only labor-intensive but also error prone. While 2D classification can categorize different states and views of helical assemblies, it can also have limitations as the filaments may not be homogenous within each 2D class (Fig. 2B-C). Additionally, the features of the 2D average images may not be clear enough to visually group them when each 2D class also represents different, unknown views and different parts of the pitch of the helix. The chep method^24, 25^ clusters the per-filament histogram of 2D class assignments to separate the different filament types. In this approach, the relative relationship of the segments along the filaments are ignored, which might cause the weak separation of the amyloid filaments^25^. Although 3D classification is the most accurate approach, it requires prior knowledge of the helical parameters and initial 3D models for each type of helical assembly, which restricts its use with experimental samples with limited prior information.

**Figure 1:**
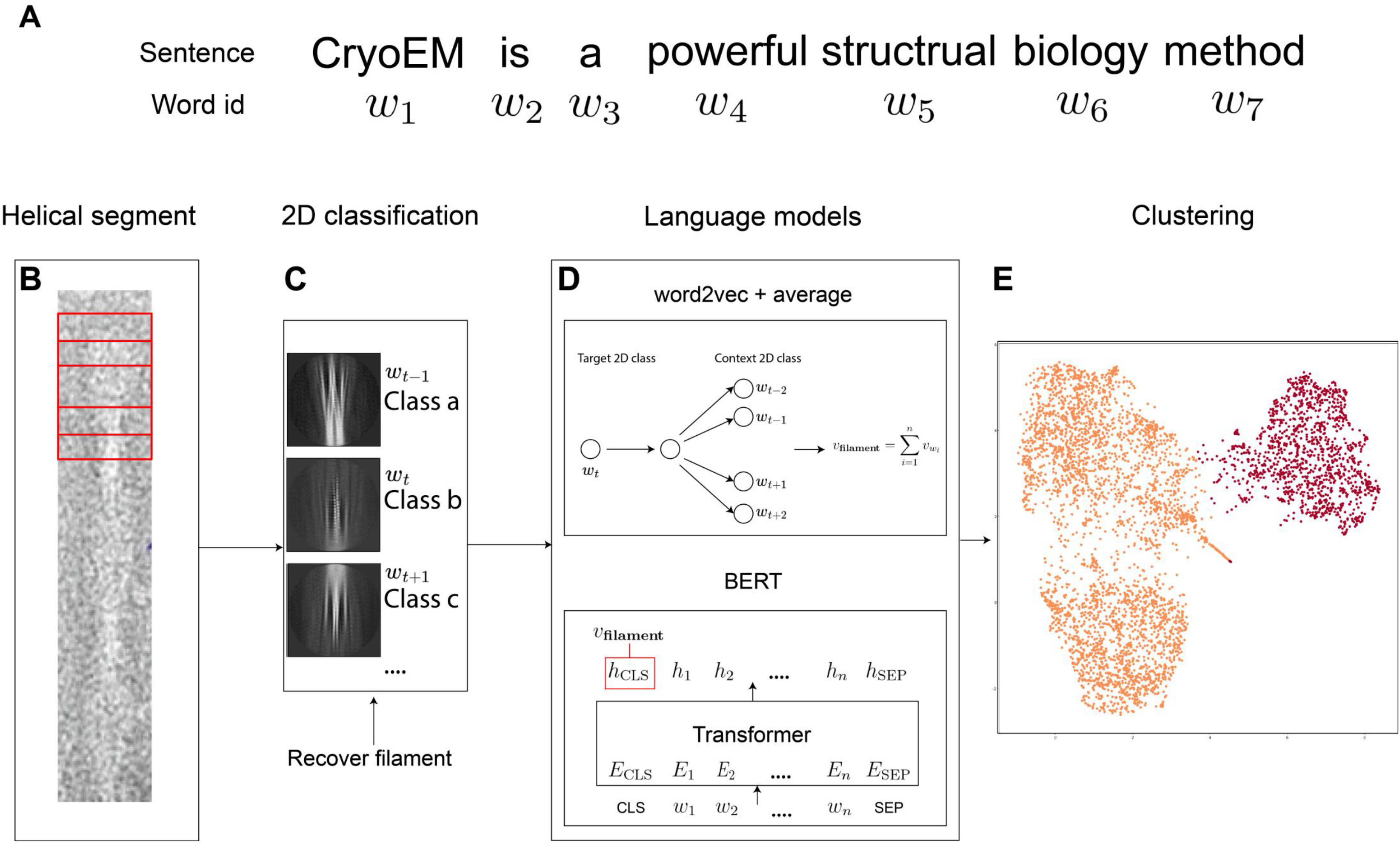
The HLM pipeline for classifying different types of helical polymer images. A. An example of tokenizing the sentence. B. Dividing the long filament into a series of short segments. C. 2D classification of the 2D segment images. If there are too many junk particles that necessitate multi-round 2D classifications and selections, an additional filament recovery process can be applied to reduce the fragmentation of filaments. The filament is converted to a series of 2D class indexes, analogous to a sentence. D. word2vec or BERT is applied to the dataset of the filament “sentences” to represent the filaments as vectors in a hyperspace. E. Classifying different types of filaments by dimension reduction and clustering of these vectors.

**Figure 2:**
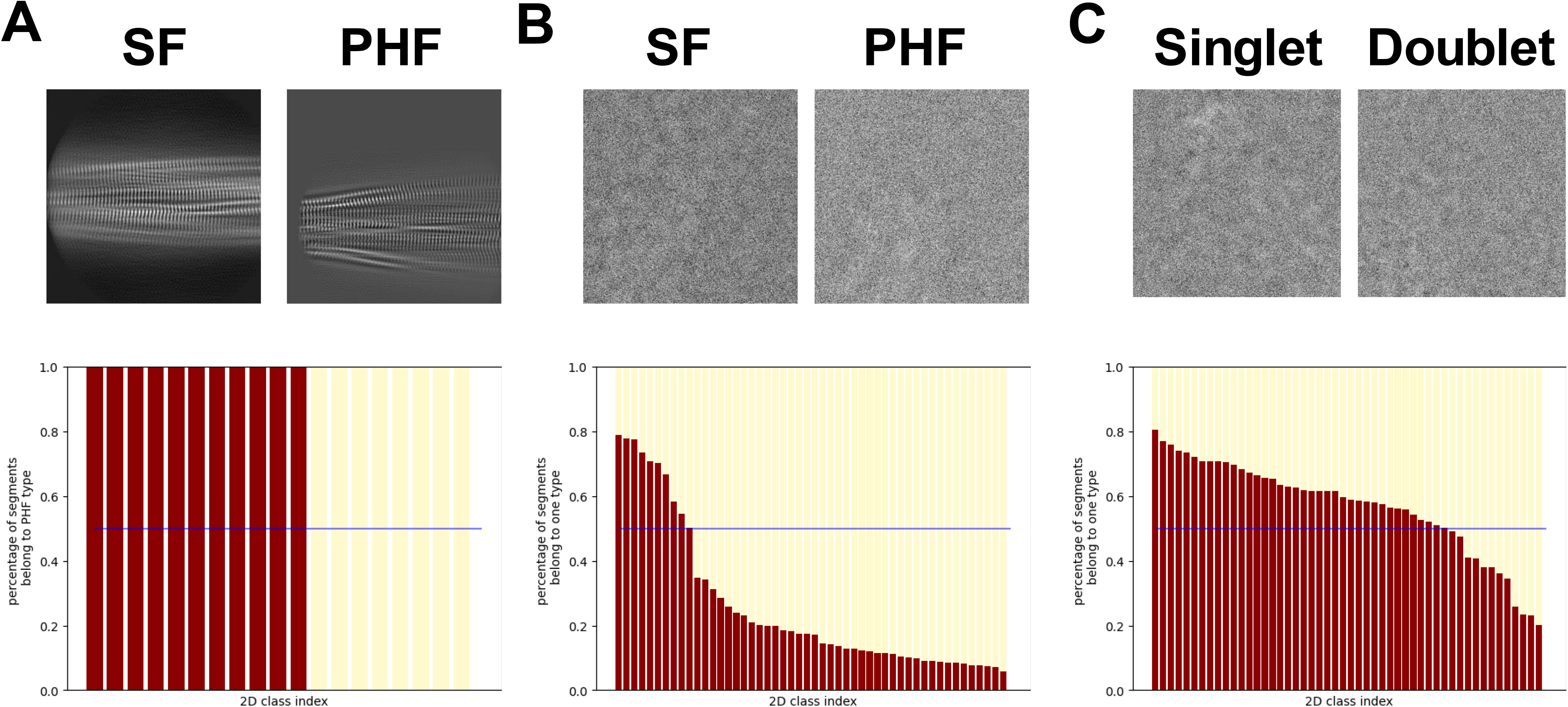
Quality of 2D classifications. The upper panel shows the example images of each test dataset. In the bar plots (lower panel), the x-axis is the 2D class index assigned by RELION, and the y-axis indicates the fraction of corresponding types of filaments. A. Simulated AD Tau images. B. EMPIAR-10230 dataset of AD Tau. C. EMPIAR-10340 dataset of CBD Tau images. For B and C, the filament type assignments included in the deposited dataset were used as the ground truth.

Here, we aim to develop a computational method to quantitatively separate the different types of helical structures in a cryo-EM dataset. After 2D classification, each of the helical segment images is assigned a 2D class index (Fig. 1B-C). These indices, when associated with the sequential position information along each filament, can be considered special sentences, with each 2D class index serving as a ‘word’ (Fig. 1A, C). In recent years, self-supervised learning of natural languages has been shown to be extremely powerful not only in the quantitative representation of words and sentences^26–28^ with successful applications such as document clustering^29^, and generation of new documents^30^, but also in biomedical research, for example, in protein informatics^31, 32^ and structural prediction^33, 34^. Thus, we explored the language model technique to classify different helical structures.

In this Helical classification with Language Models (HLM) method, we have examined two self-supervised learning methods: the classic word2vec model^26, 27^ and a recent Transformer based BERT model^28^ for helical classification (Fig. 1D). Both models have been shown to be effective in separating different types of manually picked or auto-picked filaments in simulated and experimental datasets. For more challenging datasets, we augmented the 2D classification and class selection results with filament recovery to reduce the number of fragmented filaments (e.g., sentences with missing words). In one of the validation tests, it was shown that HLM-Transformer led to the discovery of a subset of Tau filaments in a public dataset that has an extra, unreported non-proteinaceous density.

## 2. Method

### 2.1. Simulation data

The first simulation experiment involves two helical tube structures, TMV (EMD-10129)^8^ and VipAB (EMD-2699)^35^, and the second simulation experiment involves two amyloid tau structures, EMDB (EMD-0259 for the PHF type, EMD-0260 for the SF type)^1^. The 3D density maps of these test structures were projected at random angles around the helical axis and in-plane orientations using the EMAN2 software^36^. Different levels of Gaussian noise (σ=1,10) were added to test the HLM method’s robustness against noise.

### 2.2. Experimental data

Three experimental datasets of tau, EMPIAR-10243, EMPIAR-10230 and EMPIAR-10340, were used as test datasets. Three different methods were used to select the filaments. The first is to use the manually picked coordinates included in the datasets. Two auto-picking algorithms, RELION’s templated-based filament picker^37^ and the Topaz filament picker adapted for RELION^38^ were also used to test the HLM performance for different qualities of filament selections. Helical segments were then extracted from the selected filaments using RELION with the size of the extraction box according to the ones reported in the literature^39^, and the inter-box distance was set to 3 times of the rise for the EMPIAR-10230 dataset and 10% of the box size for the other two datasets.

### 2.3. 2D classification

We used the RELION unsupervised 2D classification method^39^ to cluster the helical segments into “classes” that have different views or structures. Preliminary experiments were performed to determine the optimal number of 2D classes and in-plane angular sampling angles. Our final choice was to use 50 classes, the T number equal to 2, and an in-plane angle sampling rate of 2 degrees for the experiments.

### 2.4. Filament filtering and recovery of full filaments

In the 2D classification process, we typically select the “good” 2D classes that display clear features for further processing. However, not all segments in the ignored 2D classes are necessarily low-quality helical segments (i.e., false negative), and conversely, the segments in the selected 2D classes do not guarantee that they are of high quality (i.e., false positive). Thus, we adopted a filament filtering strategy to mitigate this potential error caused by imperfect 2D classification. The full filament, with all its segments including those assigned to “bad” classes, will be accepted for further processing if the fraction of selected segments in the filament exceeds a certain threshold. Conversely, filaments, with all segments in the filaments including those assigned to “good” classes, will be discarded if they fall below the threshold. The threshold used in this study is 50%. This clean-up process is conceptually analogous to removal of fragmented sentences with missing words.

### 2.5. Word2vec model

After 2D classification, each helical filament can be represented as a sequence of 2D class indices. Hence, we can describe a single filament as

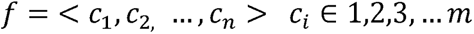

in which *m* is the number of 2D classes and *c_i_* is the 2D class index assigned to the i-th segment on the filament.

To quantify the meaning of each 2D class index, we aim to encode the 2D class indices into vectors in an embedding space, in which similar 2D classes would be closer and dissimilar classes would be far apart. Word2vec is one of the popular methods that can create word embedding based on the cooccurrence of the words in sentences^27^. To utilize the word2vec method, we would consider a filament as a sentence formed by “words” that are class indices assigned to the segments in the filament (Fig. 1A). In HLM, we implemented the word2vec model according to the word embedding pipeline developed by Mikolov et al^40^. After word2vec model has been trained to embed the 2D class indices, the filament embedding is calculated as the average of the 2D class index embeddings for the segments in the filament.

### 2.6. Transformer model

The Transformer-based models have in recent years “transformed” most machine learning tasks and become the dominant approach for a wide range of tasks, ranging from language translation and generation to image tasks^41, 42^. The same input filament “sentences” for the word2vec model are also used for the Transformer model. However, unlike real sentences, a filament with a series of 2D class indices does not have a fixed starting position due to the unknown starting axial-rotation angle of the helical polymer. Moreover, the order of a series of the 2D class indices on the filament is not as structured as a sentence. It is because the 2D class indices sometimes repeat randomly in a certain range, and the filament’s polarity is unknown. Thus, positional embedding is not used in this experiment.

We then adopted the BERT Transformer model, as described by Devlin et al^28^. The CLS token in the final hidden space embedding is used as the embedding for the filament and can be used for the downstream clustering task.

### 2.7. Dimension reduction and clustering

After obtaining the embedding for each filament using either the word2vec or the Transformer model, the dimensions of embedding vectors, which are typically set to 50, need to be reduced for visualization and clustering. We used the Uniform Manifold Approximation and Projection (UMAP) method^44^ with the default parameters.

If the separation between different clusters in the distribution is distinct enough to visually determine the number of clusters, we use the k-means clustering method with a specified number of clusters based on visual inspection^45^. Otherwise, a silhouette test is conducted first to determine the number of potential clusters^46^. All the clustering tasks were carried out using the sklearn package^47^.

### 2.8. Helical reconstruction from segments

As we are testing this method using the publicly available EMDB structures and EMPIAR datasets, the helical parameters are obtained directly from the corresponding publications. Based on the known crossover distance, the *relion_helix_inimodel2d* program is used to generate the initial model^39^. We use RELION for the 3D classification of each cluster of helical segments. The T number is equal to 8. The central z-length is 10% of the box size.

## 3. Results

To validate our HLM method, we performed multiple tests of different difficulty levels, starting with simulated datasets with clean map projections and computationally added noise, and then moving on to experimental cryo-EM datasets of *ex vivo* cross-beta amyloids with contaminants and mis-identified filaments.

### 3.1. Simulation tests

We first tested HLM with simulated datasets of two helical tubes (TMV and VipAB) computationally mixed at different noise levels (Fig. 3). When the noise level is low, both the 2D classification and HLM can effectively separate the different types of helical tubes. For datasets with higher levels of noise, the 2D classes increasingly have a mixture of multiple types of filaments (Fig. S1). However, the HLM method still shows a clear, accurate separation of the different helical tube types simulated at all the noise levels (Fig 3a-c).

**Figure 3:**
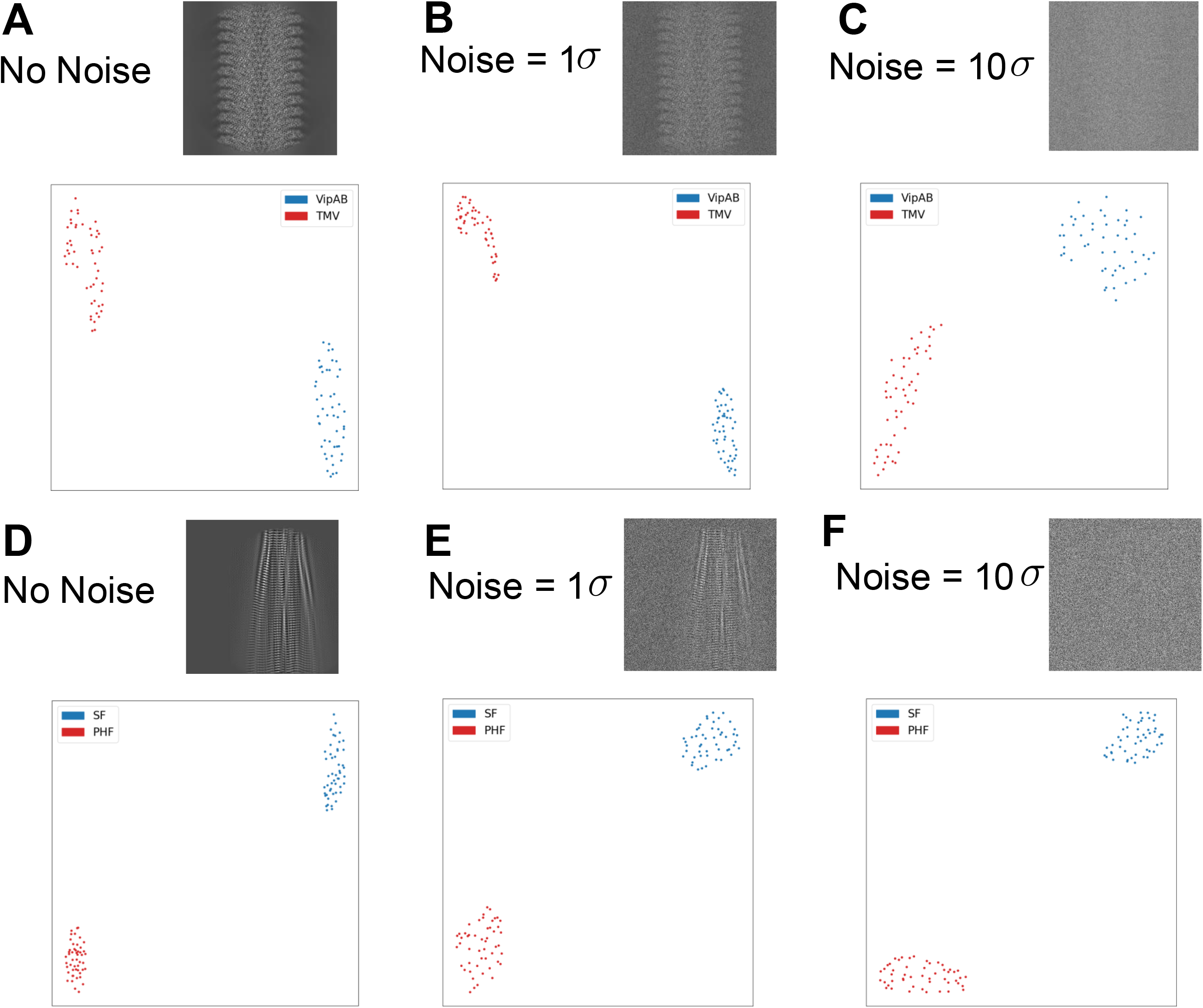
Simulation tests with HLM-word2vec. A-C: Simulated helical tube datasets with the noise level of 0, 1, 10 times of the sigma of the image. D-F: Simulated tau amyloid datasets with the noise level of 0, 1, 10 times of the sigma of the image. The distinct colors represent different types of helical polymers. The top right figure of each subplot is an example of the simulated image. In the subplots, the HLM-word2vec embedding vectors were dimension-reduced to 2d and shown as a point. Each data point represents a helical segment colored by the ground-truth type of the helical structure.

As the TMV and VipAB tubes are large (18 nm and 29 nm in diameter, 36.4 kDa and 100 kDa per nm length) and have relatively short pitch values (2.3 and 4.4 nm), we then tested HLM with simulated datasets of PHF and SF tau filaments that are much narrower (7-15 nm in diameter), have much less mass (8.9 kDa/nm length), and much longer pitch (∼80 nm). In contrast to the strong layer lines and rich information in the low resolutions (100-5Å) for protein filaments/tubes such as TMV and VipAB, the cross-beta structures of amyloids with ∼4.75 Å rise and long pitches lack layer lines and information in a large resolution range (100-5 Å) with the only prominent layer line at ∼4.75 Å corresponding to the inter-strand distance. These unique cross-beta structural features pose serious challenges in the analysis of heterogenous amyloid images. Our tests showed that the quality of 2D classification of the simulated tau filaments is indeed noticeably worse especially when the noise level increases. Encouragingly, the HLM clustering was still able to accurately separate the PHF and SF types of tau filaments into distinct clusters (Fig. 3d-f) despite the significant levels of mis-assigned 2D classes for many of the tau segments (Fig. S1d-f).

These simulation experiments suggest that the HLM-word2vec model can effectively separate the polymorphic tubes and amyloid filaments into different clusters, even in low SNR conditions where the quality of 2D classification is significantly impacted.

### 3.2. Tests with experimental cryo-EM datasets: recombinant amyloids with manually picked filaments

We then tested multiple experimental cryo-EM datasets with the first using the recombinant hairpin-induced tau dataset (EMPIAR-10243). There are several advantages to starting with this dataset. Firstly, it ensures the quality of the helical segments for processing because the coordinates and the type assignments of the filaments included in the dataset could be used as the “ground truth”. Secondly, compared to the dataset of *ex vivo* samples extracted from brain tissues, the recombinant dataset has less contaminants, which greatly improves the quality of 2D classification. Moreover, the three types of filaments in this dataset have distinct folds and helical parameters. Due to these advantages, it is anticipated that the segments within each 2D class will be homogenous in this dataset.

After 2D classifications, we apply HLM to embed each filament into a vector and cluster these vectors. Both HLM-word2vec (Fig. 4a) and HLM-BERT (Fig. S2) show similar distribution in the embedding space with four well separated major clusters. Clusters 1 and 2 have the same type of filament but have a mirrored 2D average (Fig. S3). The four HLM clusters are highly consistent with the “ground truth” assignments (Fig. 4b-c). To verify the homogeneity of each cluster, we performed a round of 3D classifications of each HLM-word2vec cluster with the helical parameters of the three types of filaments. The filaments in each cluster are homogenous, as indicated by the 3D reconstruction results (Fig. 4d-g). The results of this test demonstrate that HLM works reliably with the dataset with high-quality helical segments with clean background and few contaminants.

**Figure 4:**
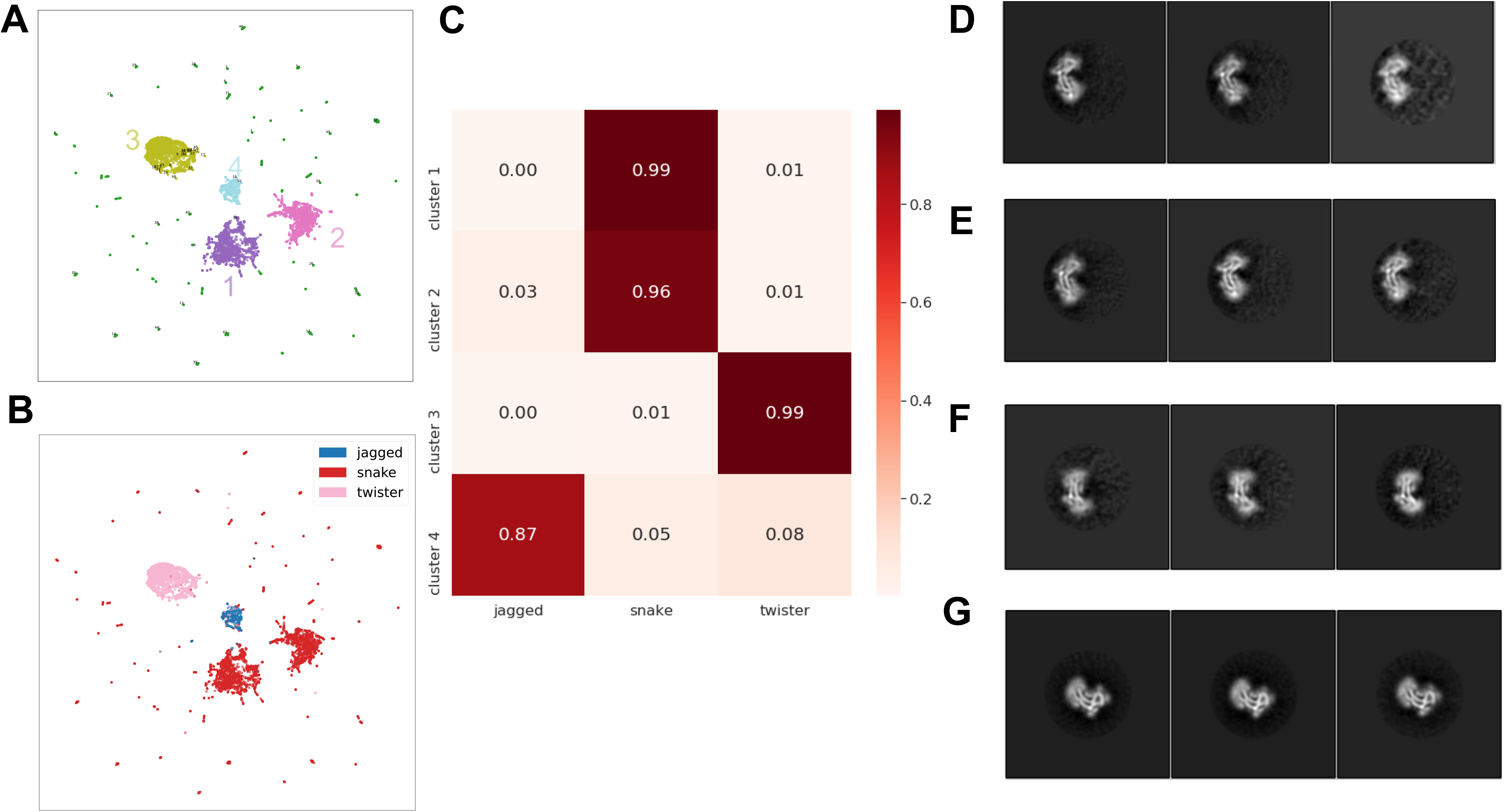
Tests with recombinant Tau dataset EMPIAR-10243. A-B: Distribution of HLM-word2vec embedding vectors where each point corresponds to one filament and is colored according to its HLM cluster assignment (A) or the “ground truth” assignments included in the deposition (B, red, snake form, pink, twister form, blue, jagged form). C: The percentage of each type of filament in each cluster according to the “ground truth” assignments. D-G: 3D classification results of the segments in each HLM cluster.

### 3.3. Tests with cryo-EM datasets: *ex vivo* sample with manually picked filaments

For experimental datasets of *ex vivo* amyloid samples extracted from brain tissues, there are often significant amounts of contaminants such as vesicles, ferritins, *etc*. in the micrographs. For instance, the AD tau dataset (EMPIAR-10230) from human brain has many ferritins that are near or over the tau filaments. As the iron-filled ferritin signal is strong, it will bias the alignment of the filaments to the ferritin density and significantly reduce the quality of 2D classification. To reduce the negative impacts of the contaminants, the filaments included in the deposited dataset were based on careful manual picking. In this HLM test, we used the manual picking coordinates included in the dataset.

There are two types of tau filaments in the EMPIAR-10230 dataset. One is the PHF type, characterized by a width varying from 70 to 150 Å, while the other, the SF type, has a more uniform width of around 100 Å. Upon application of HLM to the 2D classification results, two well-separated clusters can be observed in both HLM-word2vec (Fig. 5a) and HLM-BERT (Fig. S4) with a similar distribution in the embedding space. The filaments in the larger cluster 1 (Fig. 5a, right-side) are highly consistent (∼90%) with the PHF type of filaments as annotated in the deposition. However, the smaller cluster 2 (left-side) has ∼79% of the filaments consistent with the SF type (Fig. 5b-c). 3D reconstruction of each cluster confirmed a relatively homogenous state within each cluster (Fig. 5d-e). Despite this, there remains a significant discrepancy between cluster 2 from HLM and the manually assigned SF from the deposition. Because these two types of filaments share similar helical parameters, we can perform 3D classification with the same helical parameter and the PHF (EMD-0259) and SF (EMD-0260) maps as the initial models to verify the identity of these inconsistently assigned filaments (Fig. 5f-g). The 3D classification results have shown that there were indeed some misassignments of PHF and SF filaments in the deposition (Fig. 5f-g). We then performed 3D refinement of the segments in the HLM cluster 2 and all the manually assigned SF type segments. We found that the filaments from both subsets could reach similar resolution and 3D density map quality (Fig. 5h-i).

**Figure 5:**
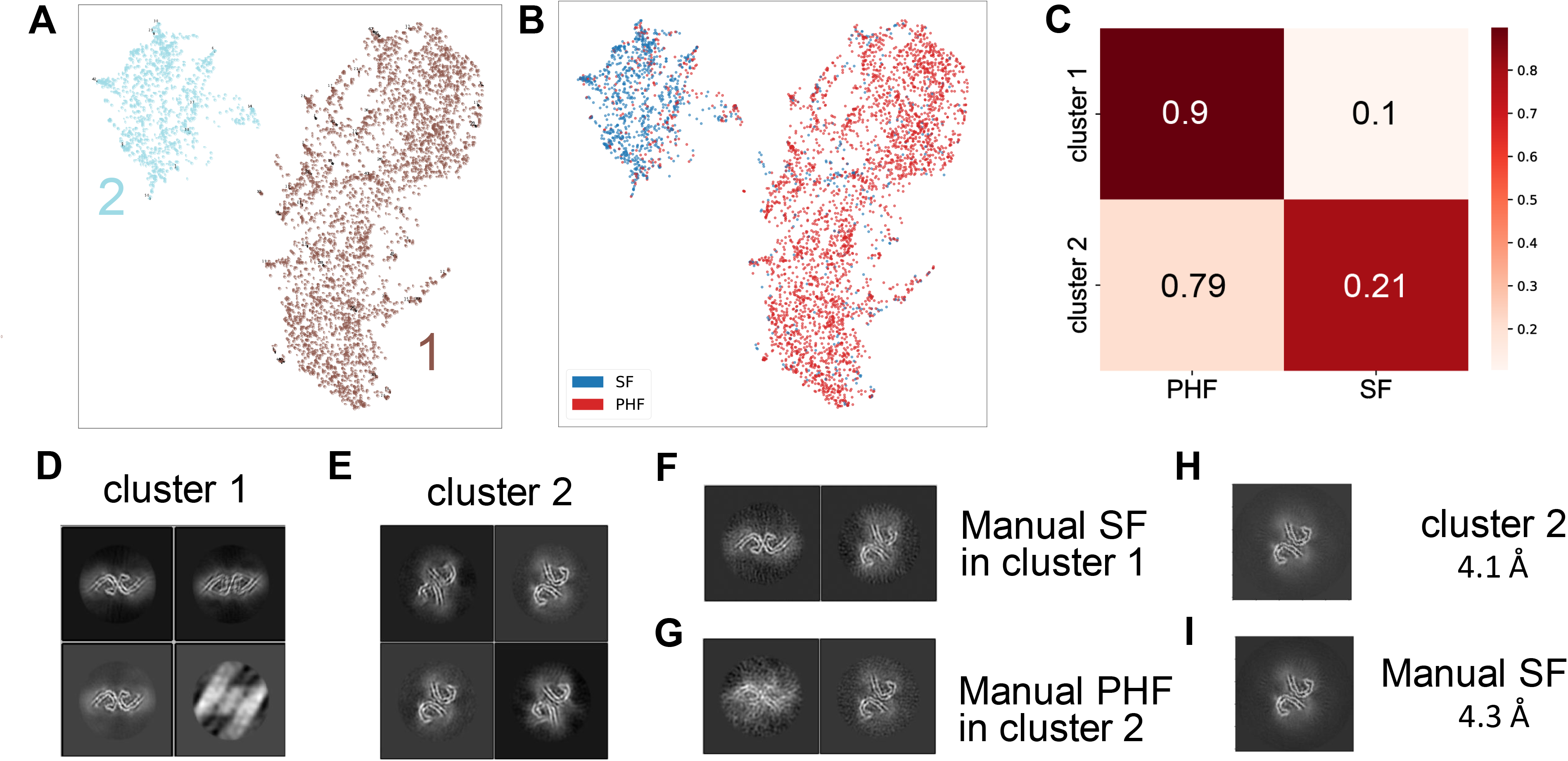
Tests with cryo-EM dataset of *ex vivo* samples with manually picked coordinates (EMPIAR-10230). A-B: Distribution of the filament embedding vectors calculated by HLM-word2vec, where each point corresponds to one filament and is colored according to its HLM cluster assignment (A) or the manual assignments (B, red for PHF, blue for SF) included in the deposition. C: The percentage of each type of filament in each cluster according to the label of the coordinates. D-E: 3D classification results of the segments in each HLM cluster. F-G: 3D classification of the inconsistently assigned filaments between the manual picks and the HLM clustering results. H: 3D refinements of the filaments in cluster 2. I: 3D refinements of the manually picked SF type filaments.

This test suggests that HLM can also be used to classify manually picked *ex vivo* helical filaments that have few contaminants. Although there are small inconsistencies between the manual assignments of filament types and the HLM clustering result, the 3D classification results have verified that the inconsistencies might be cause by some mis-assigned filament types during manual picking.

### 3.4. Tests with cryo-EM datasets: *ex vivo* sample with automatically picked filaments

Recent advances in automated particle picking have significantly improved the efficiency of cryo-EM data processing. Thus, we aim to evaluate the potential of applying the HLM method to auto-picked filaments. One of the challenges of using auto-picked filaments is that current auto-picking algorithms often include many non-filamentous densities such as ferritins, vesicles, and edges, which can negatively impact the quality of 2D classification. To address this challenge, we have experimented with the AD tau dataset (EMPIAR-10230) using two different auto-picking methods included in RELION.

The first auto-picking method is a modified version of Topaz for filament picking. The HLM procedure is applied to embed the filaments into vectors from which three major clusters are detected (Fig. 6a-b). After 3D classification of each cluster, it shows that the filaments in two of the clusters (2 and 3) are mainly PHF types while cluster 1 is mainly SF type (Fig. 6c).

**Figure 6:**
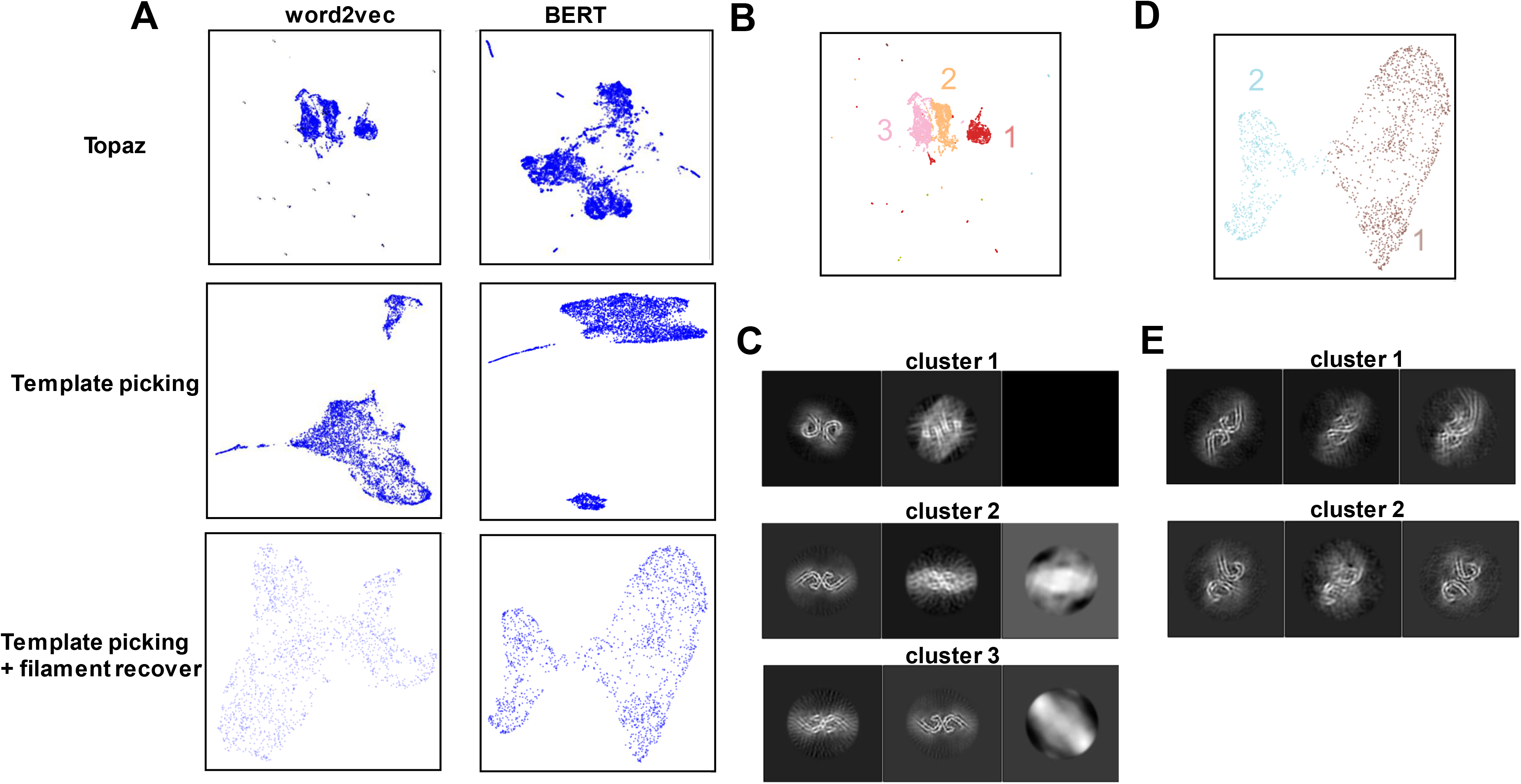
Tests with cryo-EM dataset of *ex vivo* samples using automatically picked filaments (EMPIAR-10230). A: HLM results on filaments selected by different auto-picking methods and with recovered full filaments. The bottom row was based on the 2^nd^ round of 2D classification of the main cluster in the middle row + recovered full filaments. B: The clustering result of the filament vectors using the topaz filament auto-picking method and HLM-word2vec to separate the filaments. C. 3D classification of the three clusters in B. D. The clustering result of the filament vectors from the bottom-right panel of A. E. The 3D classification of the two clusters in D.

The other auto-picking method is the template-based auto-picking with the helical mode that is implemented in RELION. Three clusters are detected after using both the HLM-word2vec and HLM-BERT (Fig. 6a, second row). The two smaller clusters seem to be mainly ferritin and junk particles (Fig. S5). The larger cluster is a mixture of both PHF and SF filaments. It seems that the HLM-BERT method can weakly separate this cluster, while no separation is noticed for the HLM-word2vec method (Fig. 6a, second row).

The HLM performance with Topaz-picked filaments is thus slightly better than that for the template-picked ones. It is probably because that the Topaz method has picked fewer non-filamentous particles than the template-based auto-picking, as fewer filaments belong to the non-filamentous 2D classes (Fig. S6).

To test if HLM performance using the template-based picking could be further improved, we also conducted an additional round of 2D classification of the main cluster consisting of a mixture of both PHF and SF filaments and recovered full-filaments to avoid fragmented filaments. The HLM method now clearly separates the filaments into two clusters (third row of Fig. 6a, Fig. 6d). The 3D classification result reveals that each cluster predominantly consists of a single type of filament (Fig. 6e). This result indicates that this HLM pipeline can be used for a dataset of auto-picked filaments in which a significant amount of junk particles are falsely included.

### 3.5. Longer segments improve the quality of 2D classification and HLM performance

We further tested HLM with another experimental dataset of *ex vivo* amyloid, CBD tau case 3 (EMPIAR-10340)^3^. There are two types of filaments in this dataset: singlets and doublets. The 2D classification results of this dataset are poor with significant mixing of the different types of filaments in the same 2D classes (Fig. 2C).

To evaluate the 2D classification and HLM performance, we tested multiple segment lengths with both HLM models (Fig. 7a). The HLM-word2vec only shows two obvious clusters with each cluster corresponding to a different type of CBD filaments when the segment length is 800 Å (Fig. 7a, left column), but only a single cluster for shorter segment lengths. On the other hand, the HLM-BERT model can separate the filaments into two marginally distinguishable clusters for all the segment lengths (Fig. 7a, right column). As a comparison, the chep method^24, 25^ can only show one cluster with all the segment lengths (Fig. S7e-g). We use k-means to separate the HLM-BERT results for the 800 Å segment length into two different clusters (Fig. 7b), and each cluster is found to correspond mainly to one type of filament (Fig. 7c).

**Figure 7:**
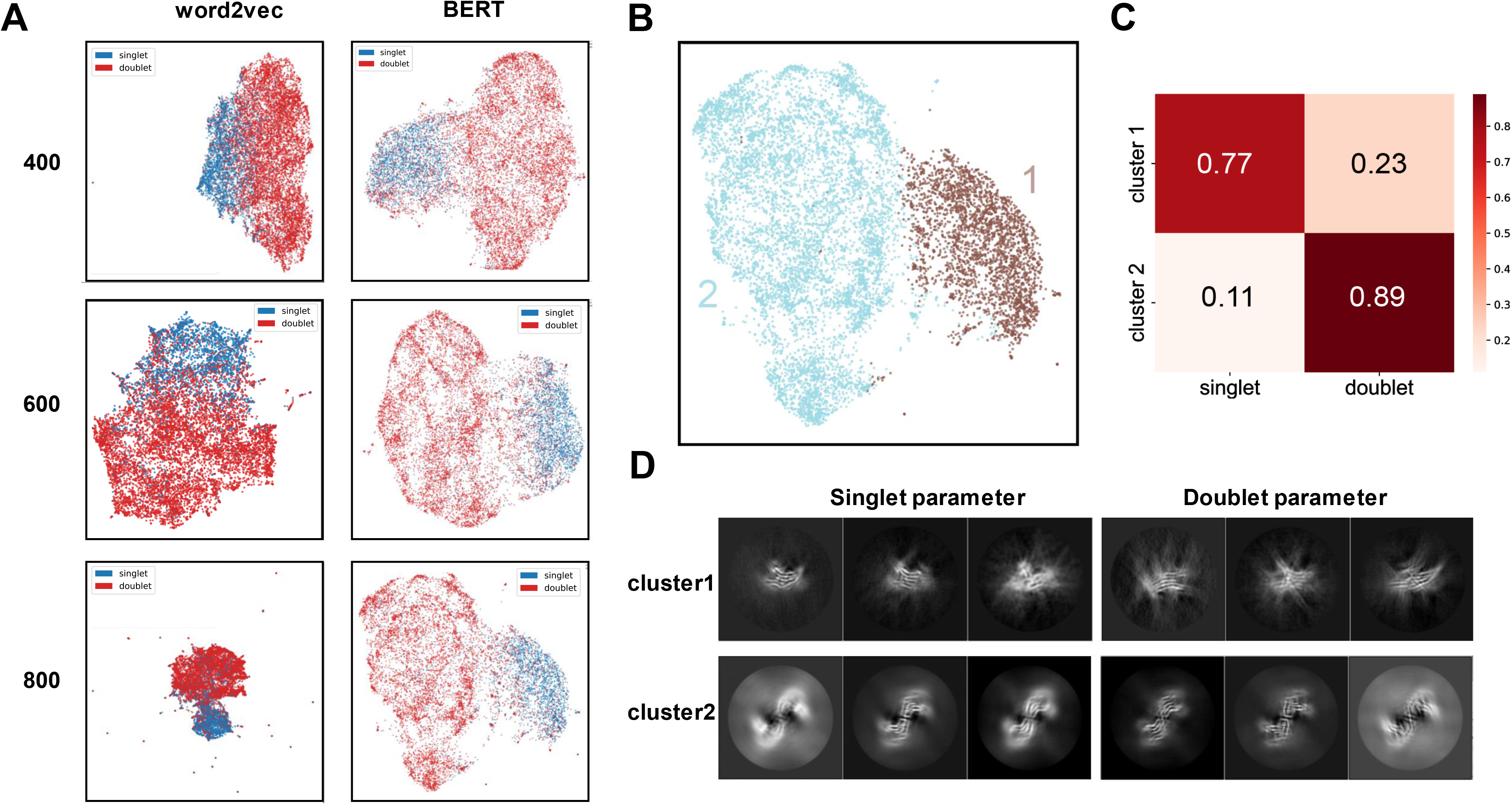
Tests with cryo-EM dataset of *ex vivo* CBD case 3 tau samples (EMPIAR-10340) with different lengths of extracted segments. A: The performance of the two HLM models on different segment lengths. B: The clustering result of the filament vectors of HLM-BERT for 800 Å segment length (bottom right panel in A). C: Confusion matrix of HLM cluster assignments and the manual type assignments (singlet/doublet) included in the deposition. D: 3D classification of each HLM cluster in B using the helical parameters of the two filament types.

We used 3D classification to investigate the identities of HLM clusters. Here, the 3D classification approach used for the AD tau dataset (EMPIAR-10230) could not be used because the two types of CBD filaments have very different helical parameters. Thus, we perform independent reconstruction of each cluster with two sets of helical parameters. The reconstructions indicate good homogeneity within each HLM cluster (Fig. 7d) by assessing the relative width of the 3D map.

This result highlights the significance of the quality of 2D classification on the performance of HLM method and the superior performance of HLM-BERT than HLM-word2vec on some challenging cases. Longer segments with more shape information resulted in better 2D classification quality and improved separation of the different filaments in the HLM results. This observation is consistent with the best practice in amyloid processing which begins with 2D classification of long, down-scaled segments^11^.

### 3.6. Discovery of a new variant of filament in the EMPIAR-10340 CBD tau dataset

The HLM is also tested on another case (case 2) in the EMPIAR-10340 dataset. The high concentration of ferritin density in the micrograph negatively impacted the quality of the 2D classification (Fig. S5). After the first round of 2D classification, HLM-BERT separates the filaments into two clusters, and one of the clusters has a high percentage of ferritin density (Fig. 8a, S6) and not further considered. The filaments in the other cluster (Fig. 8a, circled) are used for further processing, and they can be separated into two clusters by HLM-BERT (Fig. 8b); each cluster mainly consists of one type of filament (Fig. 8c,d), according to the annotated filament types included in the deposition. On the other hand, the HLM-word2vec only shows one single cluster in the embedding space (Fig. S8b.c). To verify the identity of the filaments, we performed 3D classification of each cluster with two sets of helical parameters, one for singlets and another for doublets. The homogeneity within each cluster is clear as shown by the 3D classes in each 3D classification experiment (Fig. 8e).

**Figure 8:**
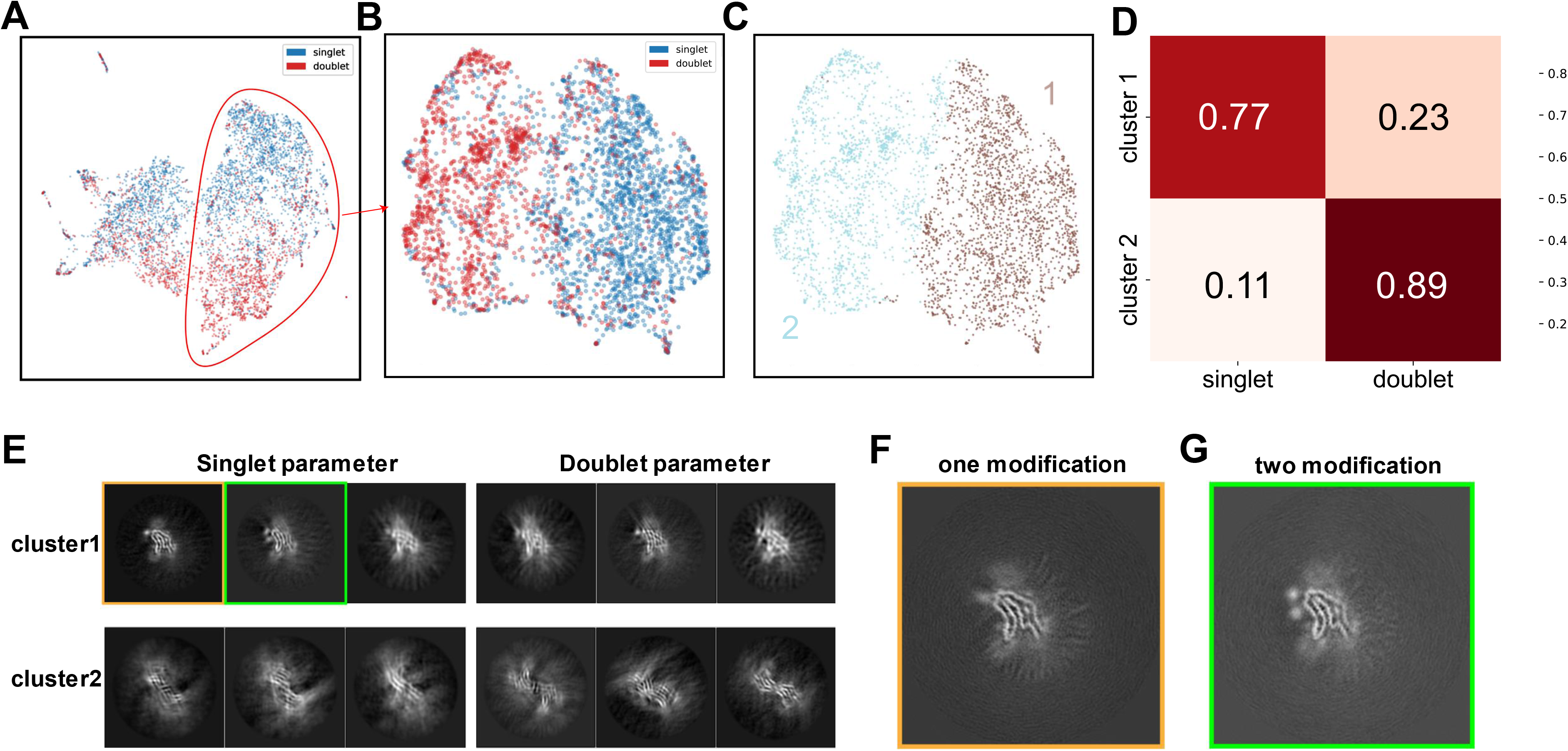
This HLM pipeline leads to the discovery of a novel CDB tau singlet filament in the EMPIAR 10340 case 2 dataset. A: Clustering of HLM-BERT embeddings of the filaments. The circled cluster with fewer ferritin contaminants is selected for further 2D classification and HLM-BERT embedding. B,C: HLM embeddings of the filaments in the circled cluster in A. The color assignment is based on the manually picked coordinates included in the deposition (B) or k-means clustering(C). D: Confusion matrix of two clustering assignments shown in B and C. E. 3D classification of the filaments within each cluster in C using the helical parameters of singlet and doublet filaments. F,G: Zoom-in view of the two types of singlet filaments (cluster 1 in C) displaying one (F) or two (G) non-proteinaceous densities.

It is interesting to note that 3D classification of singlet filaments in cluster 1 (Fig. 8c) shows not only the map similar to the previously reported singlet filament with one extra non-proteinaceous density near the tip (Fig. 8f)^48, 49^ but also a novel variant of filament with two strong non-proteinaceous densities that has not been discussed before (Fig. 8g). Further processing with a tighter mask improves the density maps to a higher resolution (Fig. S9). Cross-validation experiments with reference maps with one or two modifications show that the discovery of this novel filament is reproducible with different initial models (Fig. S10), which suggests that the difference in the assignment of filament types by HLM and the better initial model from the HLM cluster may have contributed to this discovery. Comparing the two density maps and the public CBD tau singlet type model, we can find that the new filament variant is identical to the previously reported singlet except for one extra non-proteinaceous density around the position of K340 (Fig. 8f,g) that is likely a negatively charged host factor.

## 4. Discussion

In this work, we explored the application of deep learning language models on the clustering of cryo-EM images of helical filaments. This is the first time a deep-learning language model has been used to address the heterogeneity of cryo-EM images. Validation tests on four experimental datasets from large helical tubes to amyloids with high level of contaminants have shown that our HLM method can effectively separate different types of filaments. Both HLM-word2vec and HLM-BERT model perform well in most of the cases. However, the HLM-BERT model can handle more challenging datasets. This is probably because the Transformer architecture can learn the relative weight between segments while it is a fixed value for the word2vec model, indicating that Transformer can utilize the inter-segment relationship better than the word2vec model. These results align with our finding that HLM can perform better than the chep method, where no inter-segments relationship is considered. The tests also showed that high quality filament picking is important to ensure good HLM performance as too many junk particles, for example, from auto-picking, will greatly reduce the quality of the 2D alignment and pollute the filament “sentences”. We also showed that multiple rounds of 2D classification enhanced with a full-filament recovery pipeline could effectively mitigate the challenges from the junk particles included by auto-picking. Extraction of longer segments with more low-resolution shape information for 2D classification parameters was found to be essential for some tougher datasets to ensure a good separation using the HLM method.

Interestingly, some inconsistencies have been observed between the manual assignments of filament types included in the deposition and our HLM clustering results for some datasets. Further 3D classification tests indicated that manual assignments may include some mis-assignments for noisy cryo-EM images.

Excitingly, HLM could help identify a previously unreported variant of CBD tau filament with an additional non-proteinaceous density in one of the test datasets, suggesting that HLM has the potential to uncover novel structural information in public datasets through a better separation of different types of helical filaments from which improved 3D structural analyses could be obtained. An interesting, related observation is that almost all the high-resolution amyloid structures reported in recent years only utilized 10-30% of the initially selected particles. Considering the new filament variant discovered using HLM, it will be interesting to re-analyze the “solved” datasets, especially the other >70% discarded helical segments, for potentially new structural discovery.

Though the HLM pipeline is promising, some potential improvements can be made in the future. Firstly, 2D classification results with redundant 2D classes. For example, the same particle with (ϕ:, θ, ψ) and (ϕ: + π, θ, ψ + π) Euler angles can create 2D class averages related by a mirror transform^50^. This can lead to false split of the same type of filaments into separate clusters for some datasets (Fig. 4a-c, S11), when the picked filaments are not long enough to cover the whole pitch. Secondly, the quality of the input 2D classification is often sub-optimal with different types of filaments assigned to the same class, which pollutes the filament “sentences” used by HLM. Thirdly, by generating the whole filament embedding using all segments in the filament, HLM implicitly assumes that the entire filament has the same helical symmetry and structural information. While this is true for most helical structures, this assumption may not be valid for some rare cases with rapid intra-filament heterogeneity.

In summary, our HLM method uses deep learning language models for the classification of different types of helical polymer structures. The method could help improve data analyses of helical structures by isolating the entire dataset with mixed types of filaments into a set of more homogenous sub-datasets from which a better initial model can be generated for each subset. HLM thus can be a useful data analysis step to be integrated into the cryo-EM image analysis pipeline for helical structure after 2D classification but before *ab initio* 3D reconstruction.

## Credit author statement

Daoyi Li: Software, Validation, Formal analysis, Investigation, Writing – original draft.

Wen Jiang: Software, Validation, Formal analysis, Investigation, Writing – review & editing.

## Acknowledgement

This work was supported in part by NIH grants 5U01NS110437 and 1RF1AG071177. We thank Guang Lin and Yang Ying for the discussions, and Bharath Sakshibeedu and Kadir Ozcan for proofreading the manuscript. We thank the Purdue Rosen Center of Advanced Computing for providing computational resources.

## Data availability

The HLM is implemented in Python. The source code, documentation for installation and usage is released on GitHub at https://github.com/smallelephant9516/HLM. The RELION star files for the test cases are available at https://zenodo.org/record/8153571.

## Figure Legends

**Figure S1:**
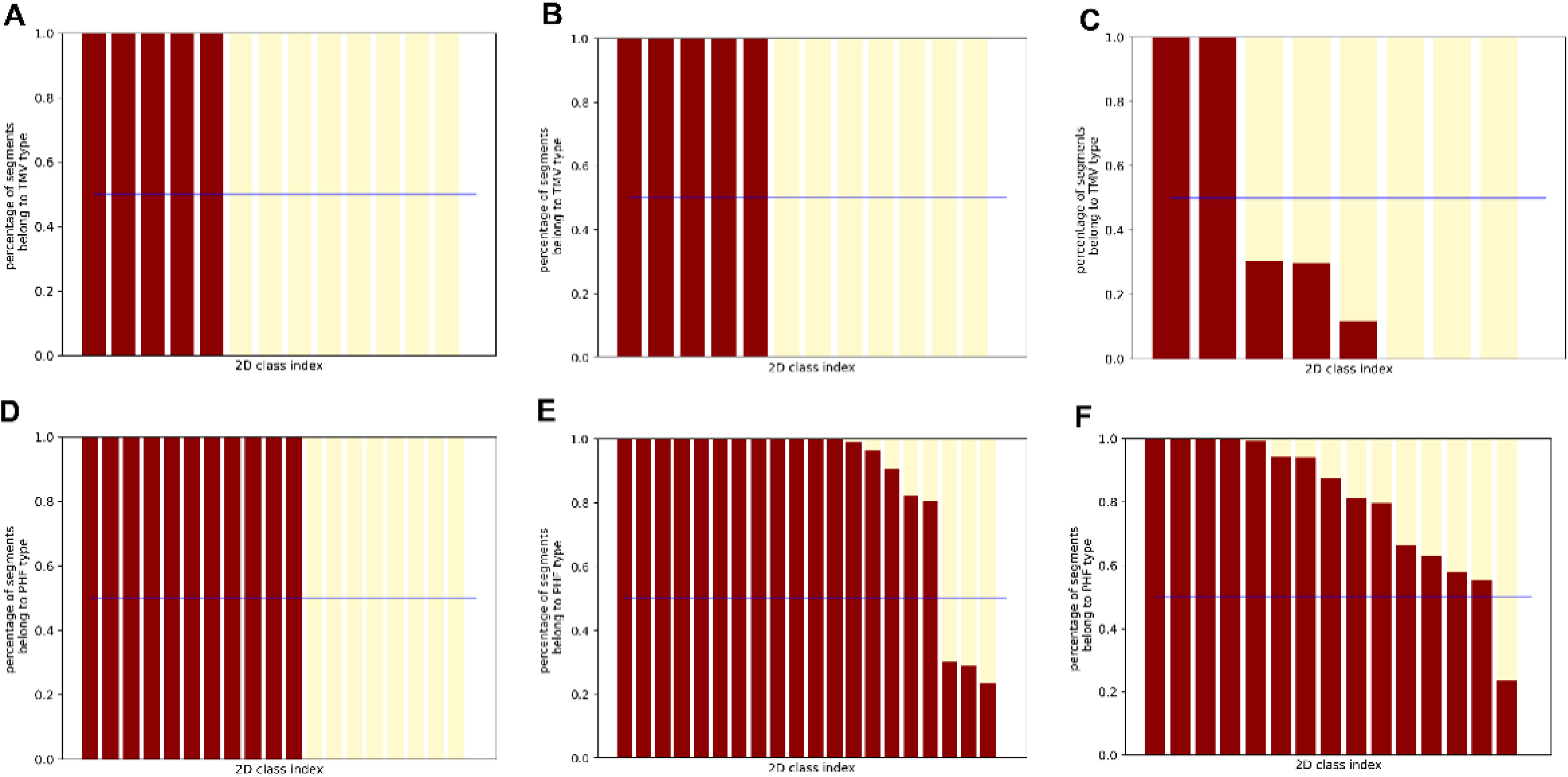
Quality of 2D classifications on the simulated dataset. In the bar plots (lower panel), the x-axis is the 2D class index assigned by RELION, and the y-axis indicates the fraction of corresponding types of filaments. A-C: The computationally mixed TMV and vipAB dataset with noise level of 0, 1 and 10 sigmas of the signal. D-F: The computationally mixed PHF tau and SF tau filament with noise level of 0, 1 and 10 sigmas of the signal.

**Figure S2:**
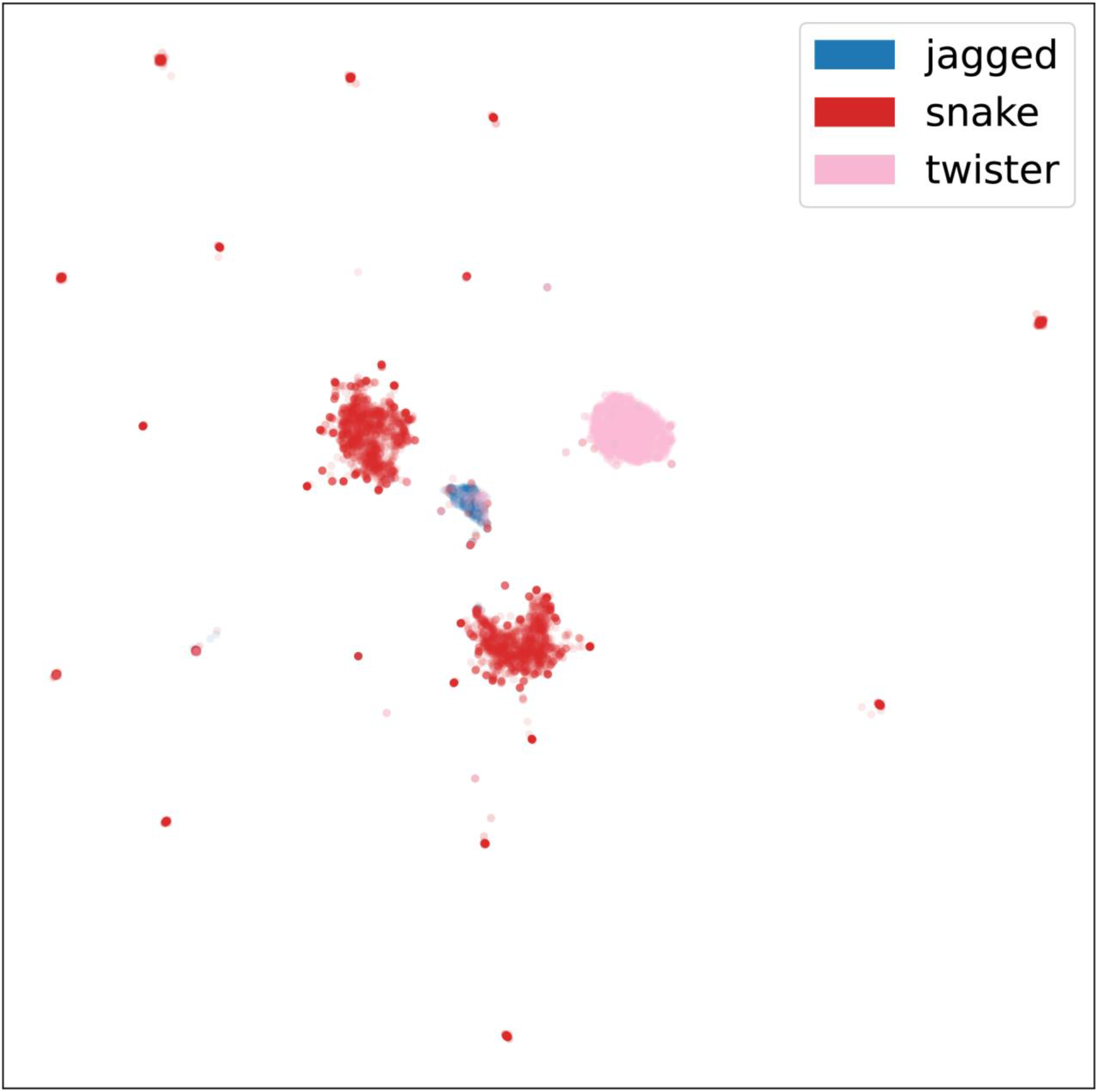
HLM-BERT on EMPIAR-10243 tau dataset. Different colors represent the manual labels for the different types of filaments.

**Figure S3:**
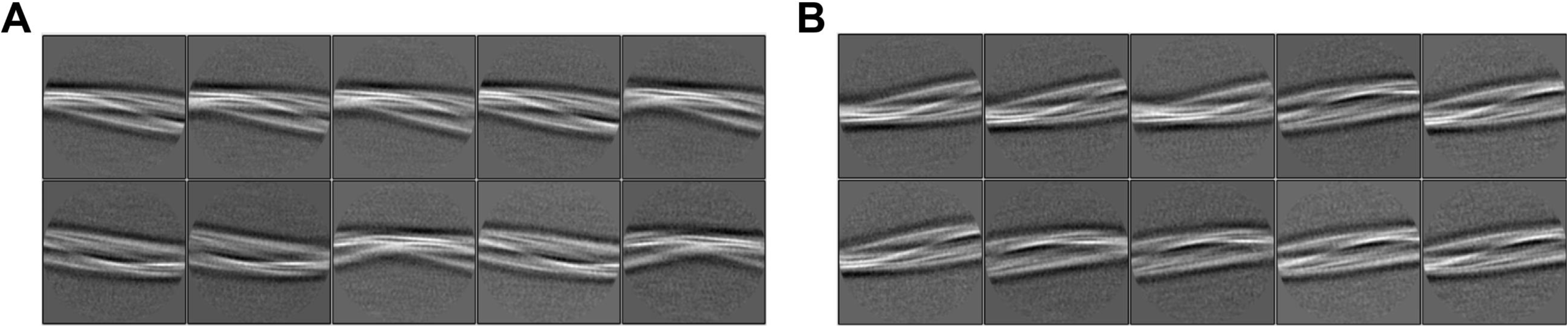
Clusters 1 and 2 in. Figure 4A **are the same type of filaments**. 2D classification of each cluster generated 2D class averages that are mirror of each other.

**Figure S4:**
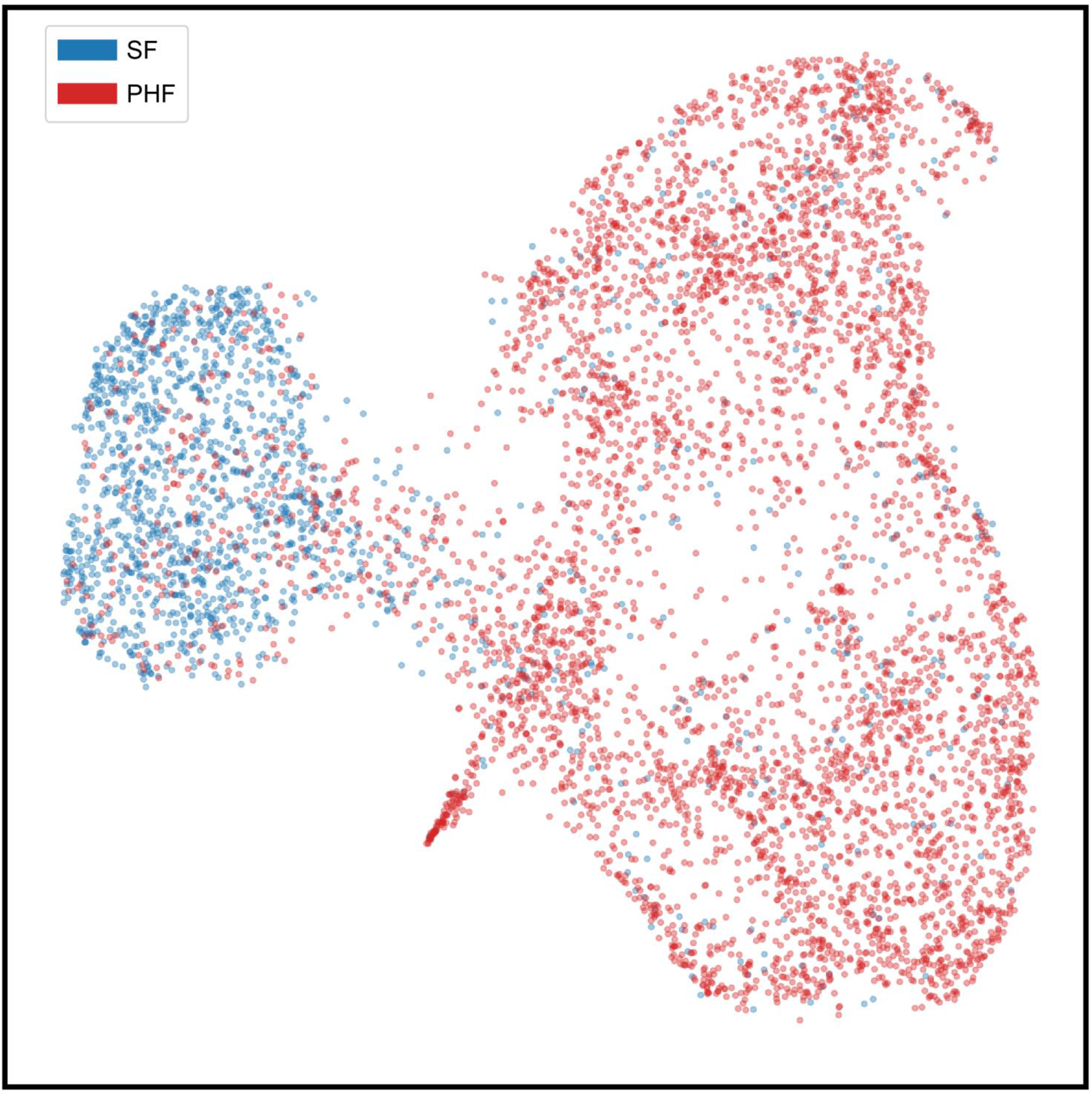
HLM-BERT on EMPIAR-10230 tau dataset using manually selected filaments included in the deposition.

**Figure S5:**
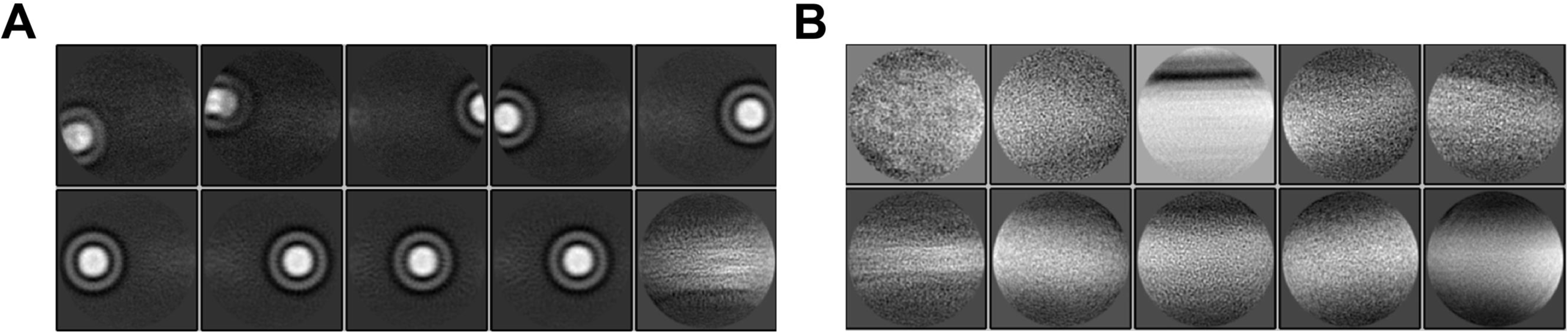
Further 2D classification of the two small clusters in Fig. 6a “Template picking” shows that these 2D clusters are mainly ferritin particles (A) and junk particles (B).

**Figure S6:**
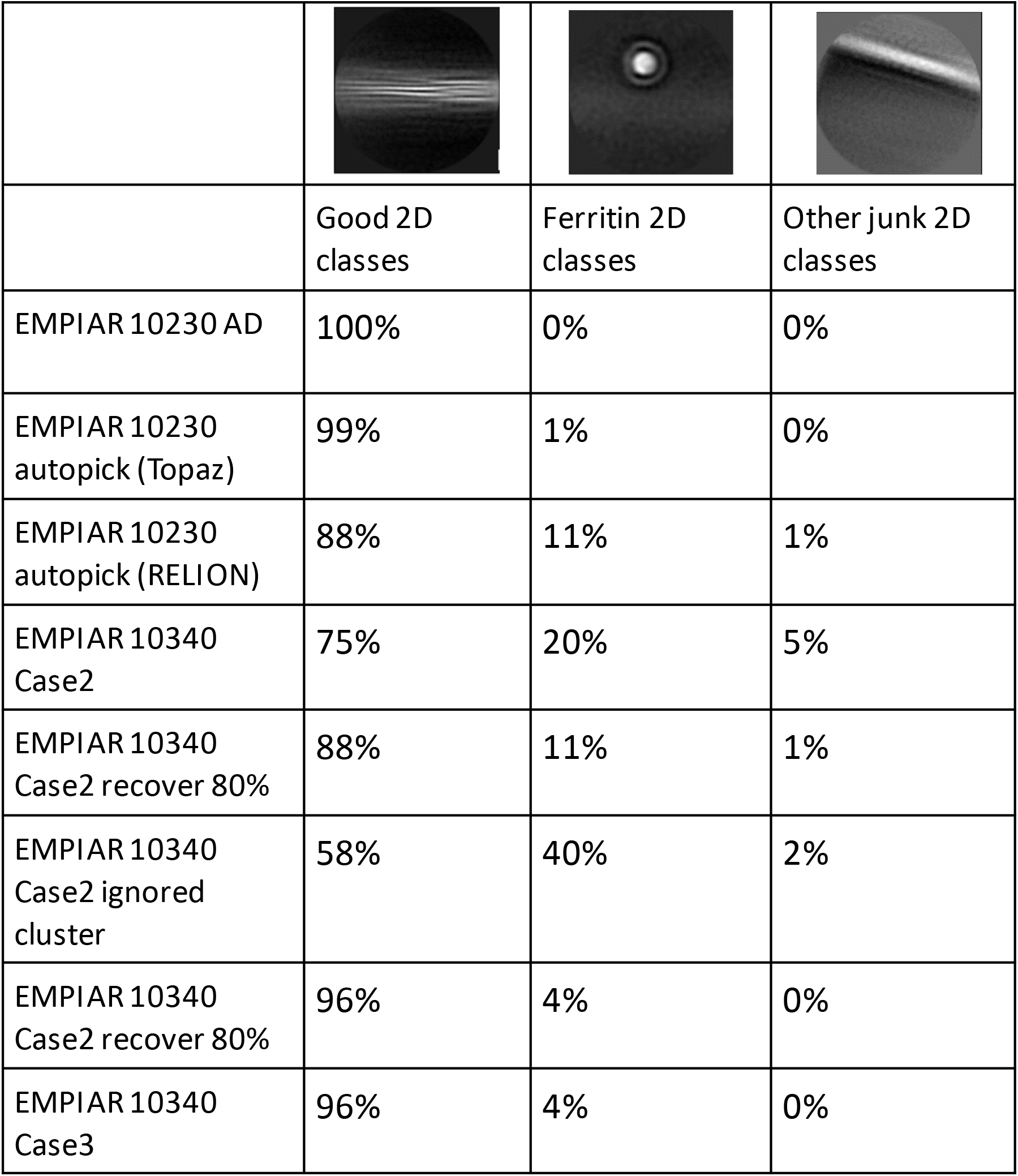
The amount of amyloid, ferritin, and junk 2D classes across the amyloid tests.

**Figure S7:**
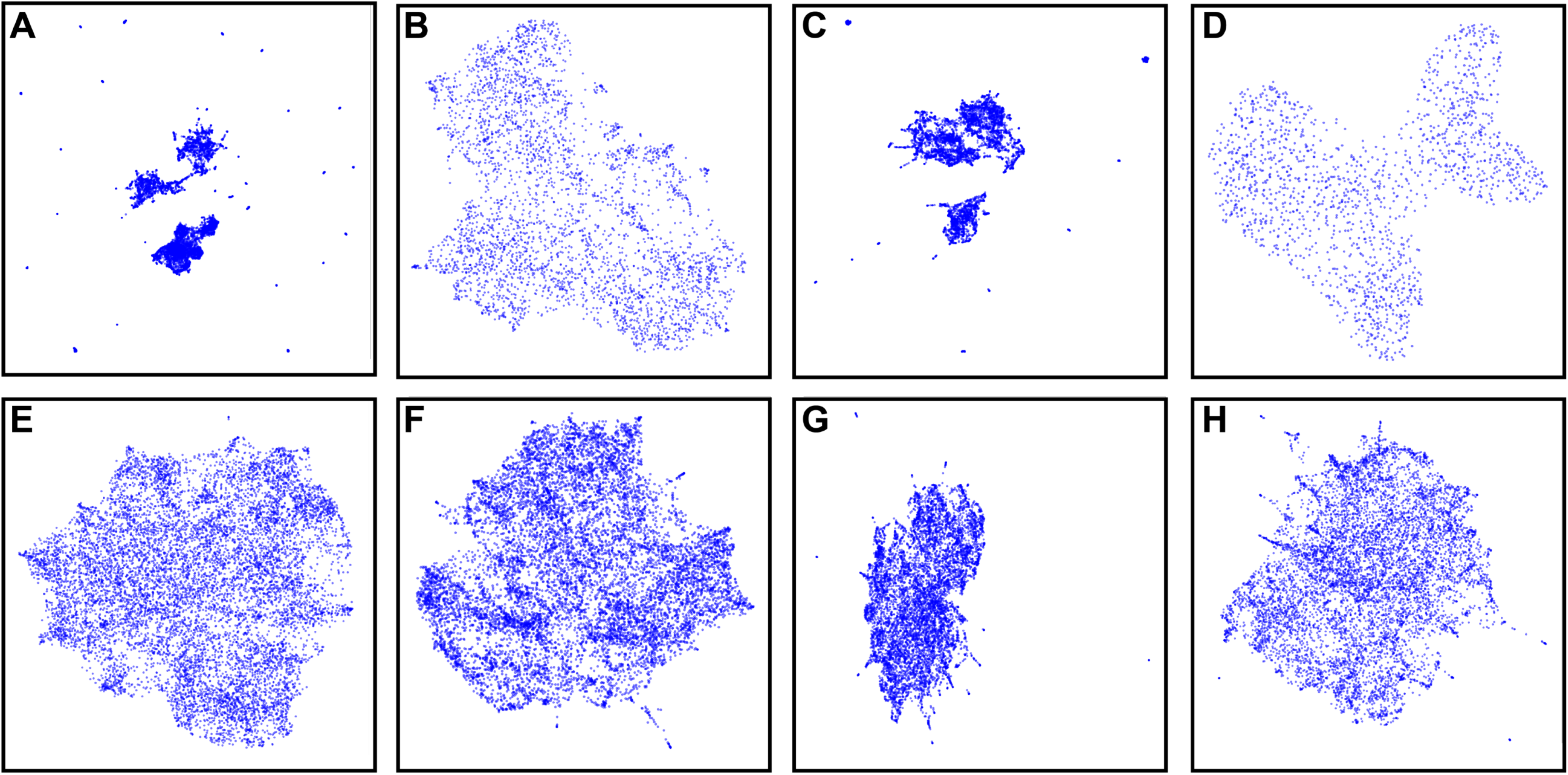
The clustering result of the chep method. A: EMPIAR 10243 dataset. B-D: EMPIAR 10230 with manually picked filaments (B), topaz auto-picked filaments (C) template picking + recovered full filaments (D). E-G: EMPIAR 10340 case 2 dataset using manually picked filaments with extracted segment lengths of 400 (E), 600 (F), and 800 (G) pixels. H: EMPIAR 10340 case 2 datasets of the circled cluster in Figure 8a.

**Figure S8:**
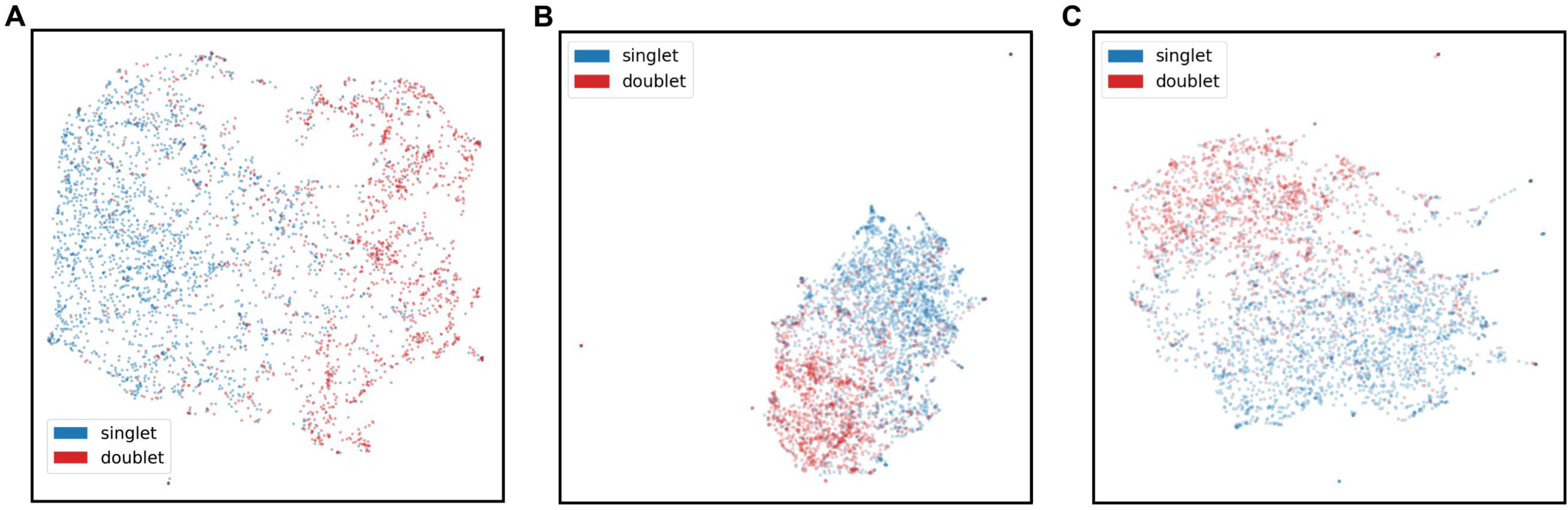
Additional HLM tests on the circled cluster in Fig. 8a for the EMPIAR-10340 case 2 dataset. A: HLM-BERT on the circled cluster without ignoring the junk particles. B: HLM-word2vec on the circled cluster without ignoring the junk particles. C: HLM-word2vec on the circled cluster after the recovering full filaments.

**Figure S9:**
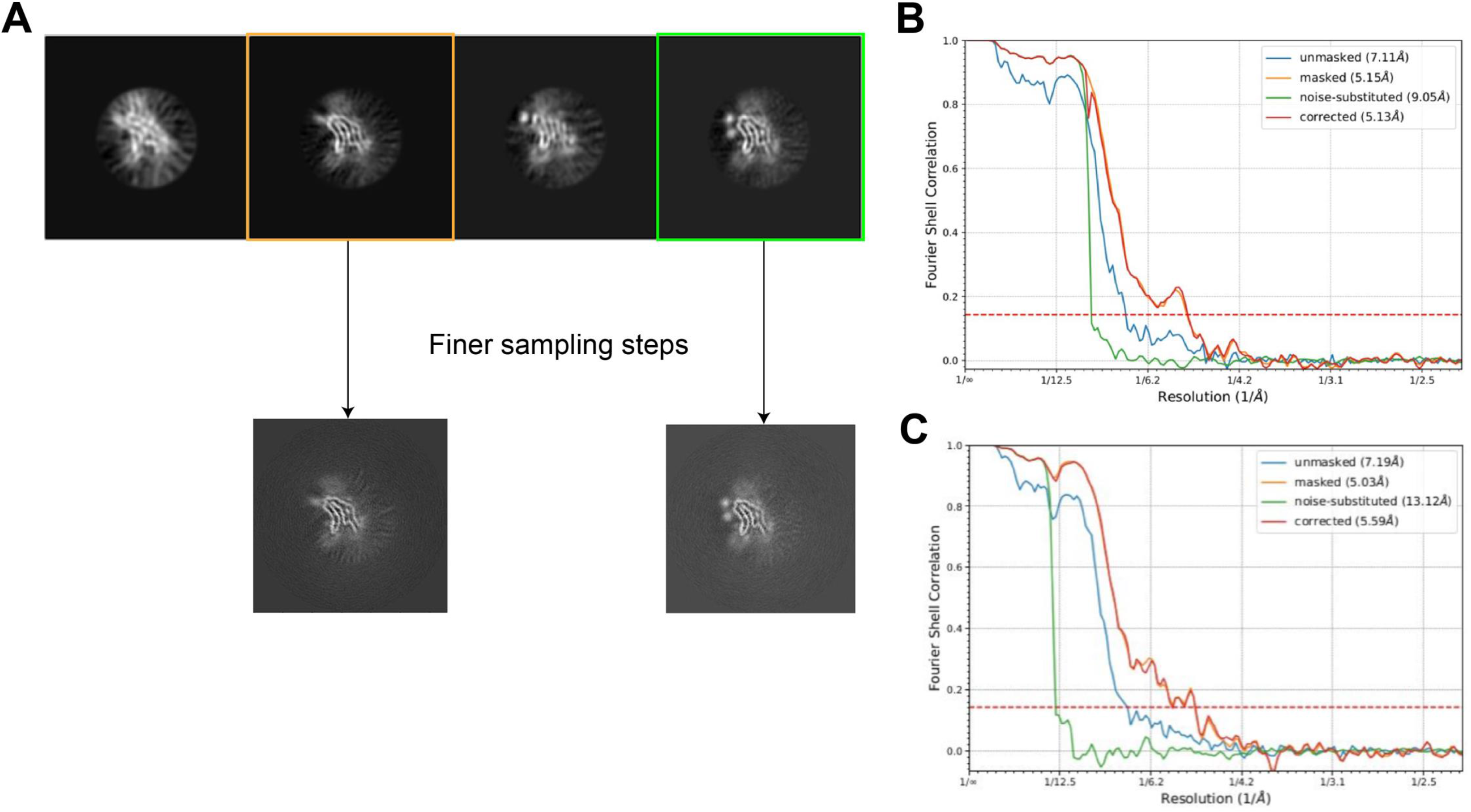
Reconstruction pipeline of the novel variant of singlet type CBD tau filament. A: The reconstruction with smaller mask size and finer angular search step size. B: FSC curve for the one modification variant of the singlet filament. C: FSC curve for the two-modification variant of the singlet filament

**Figure S10:**
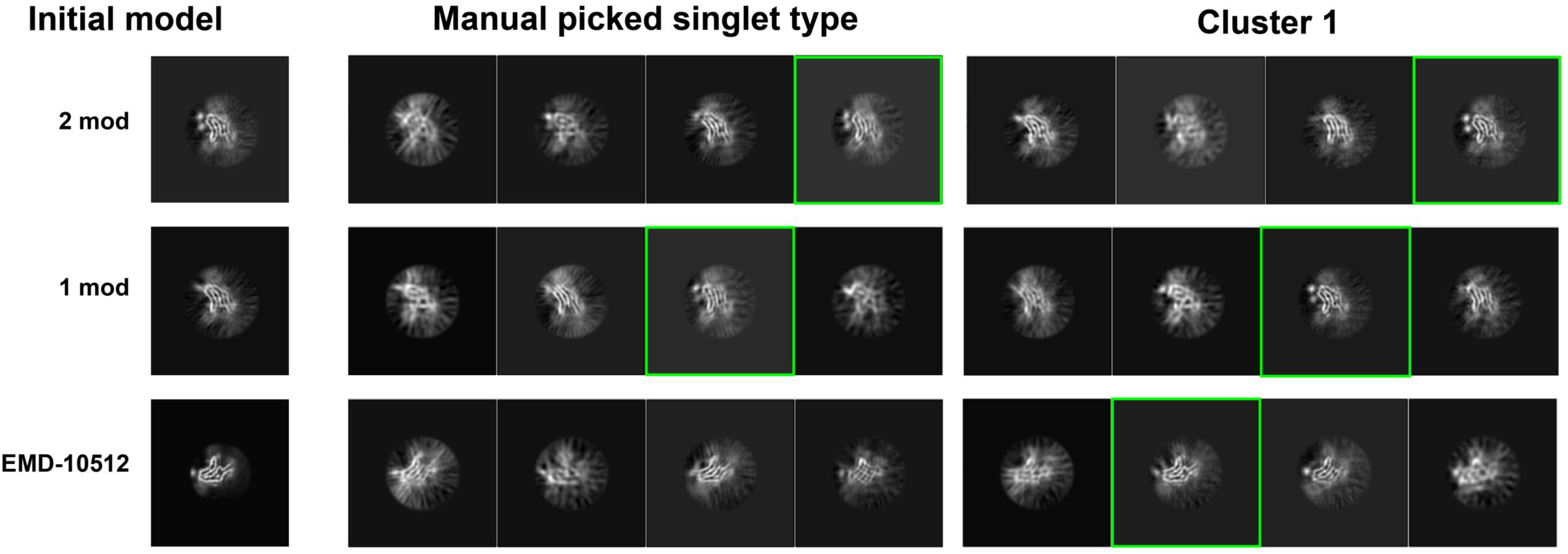
Cross-validation tests for the novel 2-modification variant of the CBD tau singlet filaments. 3D classification of the singlet type from manually picked coordinates and cluster 1of Fig. 8c with different initial models to show that the 2-modification structure is inherent to the data instead of from model bias. The quality of the reconstruction of the 2-modification variant (mark with green) in the manual picked singlet type is worse than the cluster 1. The segments from the manual picked singlet type could not recover the 2-modification variant using EMD-10512 as the initial model.

**Figure S11:**
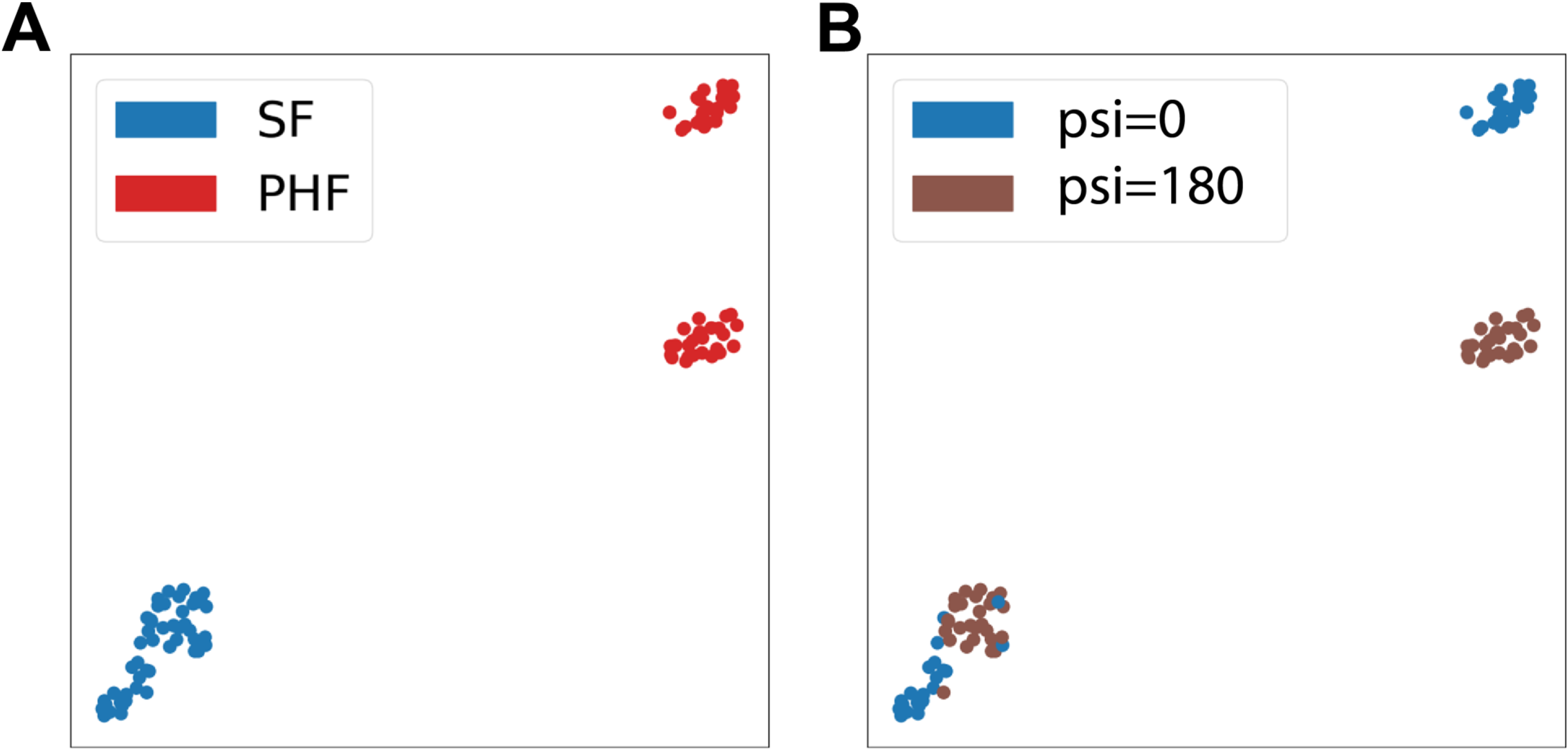
Simulation experiment with two different polarities (ψ and ψ + π) and half of the pitch with random position in a pitch. This will result in two sets of particles, one with (ϕ:, θ, ψ) and the other is (ϕ: + π, θ, ψ + π). As a result, different polarities will result in different clusters if the filament does not cover the whole pitch.

